# Zinc storage granules in the Malpighian tubules of *Drosophila melanogaster*

**DOI:** 10.1101/159558

**Authors:** Carlos Tejeda-Guzmán, Abraham Rosas-Arellano, Thomas Kroll, Samuel M. Webb, Martha Barajas-Aceves, Beatriz Osorio, Fanis Missirlis

## Abstract

Membrane transporters and sequestration mechanisms concentrate metal ions differentially into discrete subcellular microenvironments for usage in protein cofactors, signaling, storage, or excretion. Here we identify zinc storage granules as the insect’s major zinc reservoir in primary Malpighian tubule epithelial cells of *Drosophila melanogaster*. The concerted action of Adaptor Protein-3, Rab32, HOPS and BLOC complexes as well as of the white-scarlet (ABCG2-like) and ZnT35C transporters is required for zinc storage granule biogenesis. Due to similar lysosome related organelle defects, patients with Hermansky-Pudlak syndrome may lack zinc granules in beta pancreatic cells, intestinal paneth cells and presynaptic vesicles of hippocampal mossy fibers.

## INTRODUCTION

Metal ions are cofactors of enzymes (WARNER and FINNERTY 1981; KIRBY *et al*. 2008; GONZALEZ-MORALES *et al*. 2015; LLORENS *et al*. 2015). Iron and copper are required for mitochondrial respiration and cuticle formation (VILLEE 1948; ANDERSON *et al*. 2005; BINKS *et al*. 2010; KROLL *et al*. 2014), manganese for superoxide and arginine (nitrogen) turnover (DUTTAROY *et al*. 1997; SAMSON 2000), molybdenum for cysteine and methionine (sulfur) metabolism, purine and aldehyde catabolism (BOGAART and BERNINI 1981; MARELJA *et al*. 2014). The shared chemical property that turns iron, copper, manganese and molybdenum into essential cofactors of enzymes is the physicochemical stability of their ions in different oxidation states. In contrast, zinc ions do not readily change their valence and are therefore preferentially used as structural binding elements in zinc-finger transcription factors (SCHUH *et al*. 1986; REDEMANN *et al*. 1988). Alternatively, the strong Lewis acid activity of the zinc cation is utilized in proteolytic enzymes and carbonic anhydrase (WESSING *et al*. 1997; LLANO *et al*. 2000).

The development, growth, and reproduction of all animals rest on the physiological regulation of metal ions: specific protein metallation is achieved in specialized cellular compartments, facilitated by metal chaperones (LYE *et al*. 2013; QIN *et al*. 2013; SOUTHON *et al*. 2013). Such physiological regulation takes place both systemically through circulating factors secreted from specialized organs and at the cellular level through metal sensing coupled to gene and protein responses (CYERT and PHILPOTT 2013; BIRD 2015).

These systems are well understood in human iron physiology. At the systemic level, the liver senses transferrin iron saturation (i.e. sufficient iron availability), stores excess iron in ferritin, and secretes hepcidin as a response; hepcidin binds to and internalizes the iron exporter ferroportin from cell membranes, reducing iron efflux at intestinal basolateral membranes and spleen macrophages recycling iron from senescent red blood cell hemoglobin (DRAKESMITH *et al*. 2015; CAMASCHELLA *et al*. 2016; MUCKENTHALER *et al*. 2017). At the cellular level, cytosolic iron deficiency results in the stabilization of the transferrin receptor for iron uptake from circulation and in the translational inhibition of ferritin for iron storage through the action of iron regulatory proteins; the opposite effects occur under cytosolic iron overload, and these processes can be viewed as a balancing act (HENTZE *et al*. 2004; ZHANG *et al*. 2014; KUHN 2015). The similarities and differences between iron regulation in *Drosophila melanogaster* and mammals have been reviewed (MANDILARAS *et al*. 2013; TANG and ZHOU 2013). In *Drosophila*, iron availability is linked to key developmental signals, such as ecdysone synthesis (LLORENS *et al*. 2015; PALANDRI *et al*. 2015), and to processes such as the formation of epithelial septate junctions (TIKLOVA *et al*. 2010), the functionality of the circadian clock (MANDILARAS and MISSIRLIS 2012), and the induction of mitotic events (LI 2010). So far, a single iron transporter moving iron into the cytosol has been identified in flies (ORGAD *et al*. 1998; BETTEDI *et al*. 2011) and a single iron exporter has been suggested to traffic iron from the cytosol into the endoplasmic reticulum and Golgi apparatus (XIAO *et al*. 2014), where insect ferritin resides (MISSIRLIS *et al*. 2007; ROSAS-ARELLANO *et al*. 2016).

In contrast to iron, the systemic regulation of zinc homeostasis is not well understood in either human or insect biology. There is a growing appreciation of the specific, directional membrane transport functions provided by the Zrt- and Irt-like proteins (ZIPs) and Zn transporters (ZnTs) and of the metal sequestration properties of the cytosolic metallothioneins (PLUM *et al*. 2010; BABULA *et al*. 2012; KIMURA and KAMBE 2016). Zinc-responsive gene regulation is largely mediated through the Metal Transcription Factor-1 (MTF-1) (GUNTHER *et al*. 2012). The metallothionein genes are major targets of MTF-1 because the encoded proteins sequester zinc and other metals such as copper or cadmium. Nevertheless, no humoral factor has been described responding to zinc deficiency, or to zinc overload. Neither is a tissue reserve known from which zinc is mobilized to meet functional requirements under conditions of dietary deprivation. The same considerations apply to *Drosophila* zinc physiology (reviewed in (RICHARDS and BURKE 2016; XIAO and ZHOU 2016)). *Drosophila* ZIPs and ZnTs are phylogenetically conserved, with different members of each family localizing to separate subcellular compartments (LYE *et al*. 2012; DECHEN *et al*. 2015), enabling zinc absorption at the intestine (WANG *et al*. 2009; QIN *et al*. 2013; RICHARDS *et al*. 2015) and zinc excretion from the Malpighian tubules (YEPISKOPOSYAN *et al*. 2006; CHI *et al*. 2015; YIN *et al*. 2017). Cellular responses to zinc are coordinated via MTF-1 and metallothioneins (EGLI *et al*. 2003; EGLI *et al*. 2006; ATANESYAN *et al*. 2011; SIMS *et al*. 2012; QIANG *et al*. 2017). MTF-1 also regulates the ferritin subunit genes, for reasons that are unclear (YEPISKOPOSYAN *et al*. 2006; GUTIERREZ *et al*. 2010; GUTIERREZ *et al*. 2013). Little is known about the mechanism of zinc homeostasis in the organism as a whole, particularly how zinc is stored in *Drosophila*.

We came to the question of physiological zinc storage by studying *poco-zinc*, a previously identified recessive X-linked mutation that causes a threefold reduction of total body zinc accumulation in laboratory strains of *Drosophila melanogaster* (AFSHAR *et al*. 2013). By genetic mapping, we show that mutants in the *white* gene (MORGAN 1910) have a threefold reduction in zinc content. The white protein encodes an ATP binding cassette sub-family G2 (ABCG2) transporter that is best known for its function in the transport of two types of pigment precursors in the pigment granules of the eye, functioning as a dimer with either of two other members of the *Drosophila* ABCG2 protein family, brown and scarlet (DREESEN *et al*. 1988; MACKENZIE *et al*. 2000). Two types of pigment granules have been identified in wild type animals on the basis of ultrastructure morphology (NOLTE 1961; SHOUP 1966). Many eye color mutants affect enzymes of biosynthetic pathways for the brown ommochromes (WILEY and FORREST 1981) and bright red drosopterins (KIM *et al*. 2013), but a subset, known as transport mutants (SULLIVAN and SULLIVAN 1975), affect the formation of the pigment granules per se. Amongst these transport mutants, we have also analyzed total body zinc accumulation in the adaptin protein complex-3 (ap-3) mutants *garnet* (*g*) (OOI *et al*. 1997), *carmine* (*cm*) (MULLINS *et al*. 1999), *ruby* (*rb*) (KRETZSCHMAR *et al*. 2000) and *orange* (*or*) (MULLINS *et al*. 2000), in *lightoid* (*ltd*) and *claret* (*ca*) that encode for Rab32 and its Guanine Exchange Factor (MA *et al*. 2004), in *pink* (*p*), which encodes for the Hermansky-Pudlak syndrome 5 homologue (FALCON-PEREZ *et al*. 2007; SYRZYCKA *et al*. 2007) and in *light* (*lt*), encoding for the VPS41 HOPS complex homologue (WARNER *et al.*) Collectively, all these proteins are required for the biogenesis of lysosome related organelles (LROs) – specialized low-pH subcellular compartments that accumulate a variety of complex metabolites (LLOYD *et al*. 1998; KRAMER 2002; DELL'ANGELICA 2009; CHELI *et al*. 2010; HARRIS *et al*. 2011). Here we describe a LRO in the Malpighian tubules of *Drosophila* that forms the major physiological zinc storage site in this animal. This zinc storage granule concentrates the entire chelatable pool of total body zinc in flies, and is distinct from the previously described riboflavin-containing granules that give the wild type tubule its characteristic yellow-orange color (NICKLA 1972).

## MATERIALS AND METHODS

### *Drosophila melanogaster* stocks

In this study, *w*^*^ and *w*^+^ refer to isogenic stocks generated in the laboratory using the *w*^*^ mutant and the *Tan3* wild type strains, respectively (AFSHAR *et al*. 2013). First, single crosses between *w*^*^ siblings were set for 20 generations. A single *Tan3* male was then crossed to a *w*^*^ isogenic female. For 20 further generations, a *w*^*^ male (always taken from the *w*^*^ isogenic stock) was backcrossed to a *w*^*^/w+ female. Finally, a *w^+^* male from the heterozygous mothers was backcrossed to a *w^*^/w+* female to re-establish the isogenic *w*^+^ stock.

A new allele of *st^e01330^* resulting from a piggy-Bac insertion into the open reading frame of the *st* gene (THIBAULT *et al*. 2004) was crossed into the *w*^+^ background and used in this work. All strains were obtained from the Bloomington Drosophila Stock Center and are listed along with the respective stock numbers (Table 1) except for X-chromosome meiotic recombination mapping stocks *cm^1^*, *m^74f^*, *sd^1^*, *os^s^* and *w^a^*, *cv^1^*, *t^1^* corresponding to #1282 and #121, respectively, and *y^1^*, *w^67c23^*; *ZnT35C^MI07746-GFSTF.1^/SM6a*, a GFP protein-trap line (NAGARKAR-JAISWAL *et al*. 2015), corresponding to #59419. The latter chromosome was also introduced into the *w*^+^ background. All flies were fed on molasses and yeast and kept at 25°C (REMPOULAKIS *et al*. 2014).

**Table 1.**
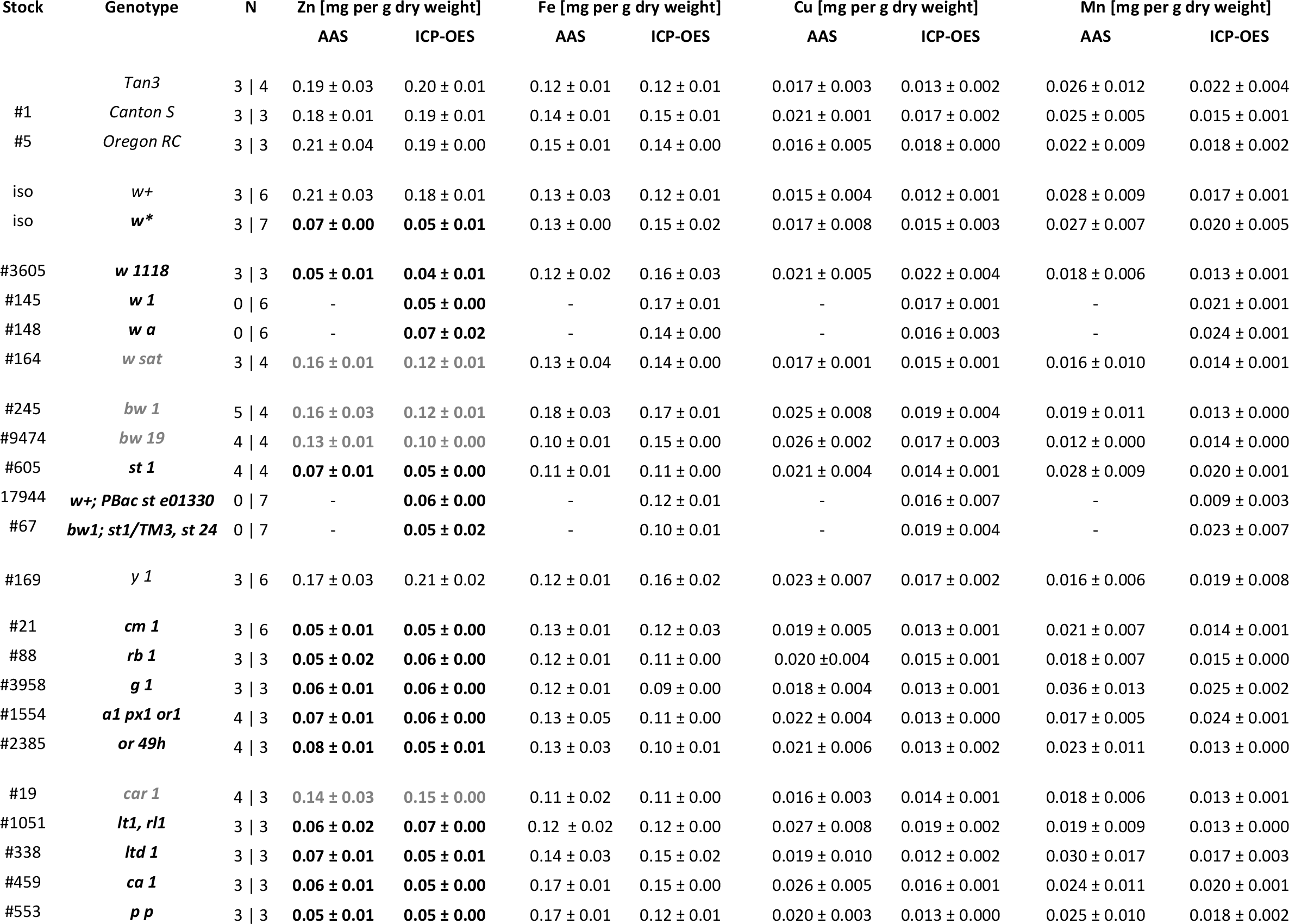
Metal determinations (mean ± standard deviation) by AAS and ICP-OES in different strains of *Drosophili melanogaster*. Bloomington stock center numbers are provided in the first column. Low zinc genotypes are shown ii bold font; genotypes with intermediate zinc are shown in grey. N corresponds to number of biological replicates respectively.

### Metal measurements

Both flame atomic absorption spectrometry (AAS) and inductively coupled plasma optic emission spectrometry (ICP-OES) were used for metal determinations. Adult fruit flies, 4-8 days old, of mixed sex were collected and stored at −80°C. They were freeze-dried for 8 hours to remove water. For the experiments with AAS 200 mg of dry sample was digested with metal-free nitric acid (Fluka) at 60°C for 48 h, whereas for ICP-OES, 20 mg of dry sample was digested at 200°C for 15 min in closed vessels of MARS6 microwave digestion system (CEM). Total Zn, Fe, Mn and Cu concentrations were measured against calibration curves and a digestion blank in the Avanta M System 300 GF 3000 AAS and the PerkinElmer Optima™ 8300 ICP-OES instruments, respectively.

### Confocal fluorescence microscopy

Malpighian tubules were dissected from adult female flies in phosphate saline buffer (PBS): 130 mM NaCl, 7 mM Na_2_HPO_4_, 3 mM NaH_2_PO_4_ (pH 7.0). The tissue was fixed with ice-cold methanol for 5 minutes and rinsed three times for three min with PBS. Fluozin-3^AM^ (Invitrogen) was dissolved in DMSO at 5 mM, stored in frozen aliquots (GROTH *et al*. 2013) and protected against exposure to direct light at all times. For each experiment, a fresh aliquot was diluted at 2.5 μM in PBS containing 0.02 *%* Triton-X100 and 0.001 % Tween20. The fixed tissues were incubated with the Fluozin-3^AM^ solution for 45 min at 38°C in a humid heat chamber. After three washes with PBS the tissues were carefully mounted in Vecta Shield with DAPI (Vector H-1200) and observed without delay under a TCS SP8 Leica confocal system coupled to a DMI6000 inverted microscope. Methanol-fixation procedure and corresponding mounting was also used for direct visualization of the ZnT35C^GFP^ construct.

### Synchrotron X-ray fluorescence microscopy

Malpighian tubules were dissected from adult female flies in PBS, washed three times with 0.1 M ammonium acetate (JONES *et al*. 2015), placed on microscope slide coverslips (Thermo Scientific™ Nunc™ Thermanox™) and air-dried at 4°C. X-ray fluorescence images were collected at the Stanford Synchrotron Radiation Lightsource using beam line 2–3. The incident x-ray energy was set to 11 keV using a Si (111) double crystal monochromator with the storage ring Stanford Positron Electron Accelerating Ring containing 500 mA at 3.0 GeV. The fluorescence lines of the elements of interest, as well as the intensity of the total scattered X-rays, were monitored using a silicon drift Vortex detector (SII NanoTechnology USA Inc.) mounted at 90 degrees to the incident beam. Photon processing was accomplished with Xpress3 signal processing electronics (Quantum Detectors, UK). In addition to these regions of interest, the entire fluorescence spectrum was also collected at each data point. The microfocused beam of 3x3 microns was provided by an Rh-coated Kirkpatrick-Baez mirror pair (Xradia Inc.). The incident and transmitted x-ray intensities were measured with nitrogen-filled ion chambers. Samples were mounted at 45 degree to the incident x-ray beam and were spatially rastered in the microbeam using a Newport VP-25XA-XYZ stage. Beam exposure was 100 ms per pixel. Fluorescence signals were normalized against the incident X-ray beam intensity to take into account its fluctuations. Data analysis was performed using the MicroAnalysis Toolkit computer program (WEBB 2011). No smoothing or related data manipulations were performed.

### Data and Reagent availability

Raw data for metal determinations and the synchrotron spectra are available upon request. The same applies to fly strains not available in the Bloomington Stock Center.

## RESULTS

### Mapping the mutant that caused three-fold reduction in total body zinc

We refer to the X-linked recessive mutant with 3-fold reduction in total zinc accumulation as *poco-zinc* (AFSHAR *et al*. 2013). The X-chromosome meiotic recombination mapping stock *cm^1^*, *m^74f^*, *sd^1^*, *os^s^* accumulated 0.07 mg zinc per g dry weight, suggesting that it also carried the *poco-zinc* allele. Flies from this stock were crossed to counterparts from wild type *Tan3*, accumulating 0.20 mg Zn per g dry weight. A total of 121 recombinants were established arising from single or double crossovers between the parental chromosomes and were screened for the presence of *poco-zinc*. The *cm* mutant was present in 48 recombinants, all of which also carried *poco-zinc*, whereas in the remaining 73 recombinants neither *poco-zinc* nor *cm* was present (Figure 1A). To estimate the distance between *poco-zinc* and *cm*, we generated a new recombinant *y^1^*, *cm^1^* stock (low in zinc) and outcrossed it with wild type flies. We were unable to dissociate *poco-zinc* from cm: 259 single *cm^1^* mutants derived from this cross segregated with *poco-zinc*, whereas 221 single *y^1^* mutants were normal (Figure 1B). To confirm the proximity between *poco-zinc* and *cm*, we also used the strain *w^a^*, *cv^1^*, *t*^1^. Surprisingly, all recombinants carrying the *w^a^* allele (irrespectively of whether they carried along either *cv^1^*, or *t^1^*, or none of the two) were associated with *poco-zinc*, and, conversely, all *w*^+^ flies were normal. Since *cv^1^* lies between *w* and *cm*, the findings pointed to a new chromosomal location for *poco zinc*, this time in the vicinity of *w* and distant from *cm* (Figure 1C).

**Figure 1.**
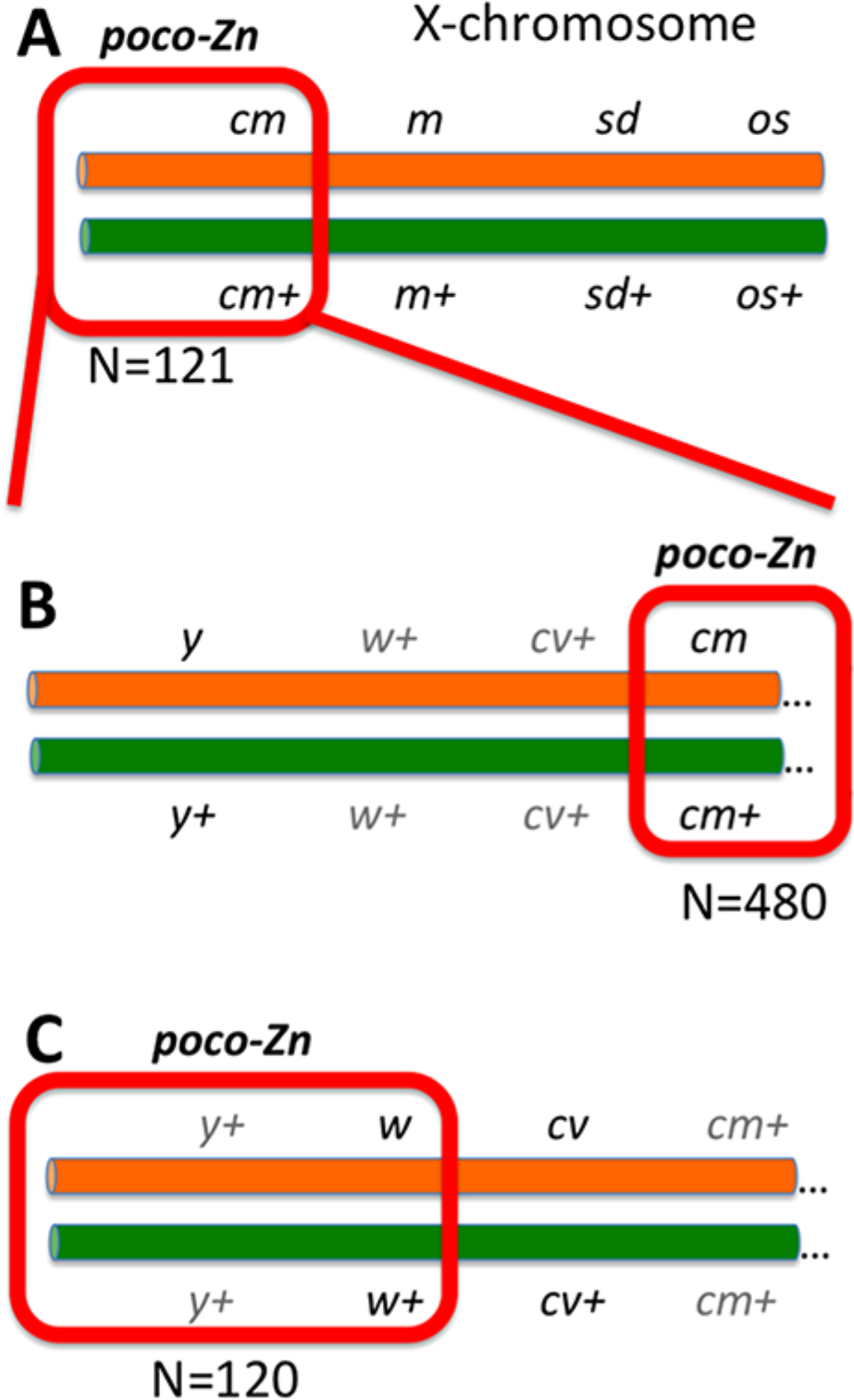
Meiotic recombination mapping strategy for *poco-zinc*. The wild type Tan3 chromosome is represented in green, whereas the mapping stock chromosome, carrying recessive alleles with visible phenotypes, in orange. (A) The first set of recombinant analysis situated the *poco-zinc* allele on the left part of the X-chromosome as it segregated 100% together with the *cm* gene. (B) Efforts to dissociate *poco-zinc* from *cm* were not successful, suggesting that *poco-zinc* is tightly linked (or identical) to the *cm* gene. (C) Efforts to map *poco-zinc* using a different mapping stock resulted in joint segregation of *poco-zinc* together with the *w* gene and far away from the *cm* gene. N depicts the total number of recombinants analyzed.

The *cm* gene encodes for the μ3 subunit of the AP-3 complex (MULLINS *et al*. 1999; RODRIGUEZ-FERNANDEZ and DELL'ANGELICA 2015). In mammalian cells, the AP-3 complex is required for the formation of LROs, organelles known to accumulate zinc (KANTHETI *et al*. 1998; SALAZAR *et al*. 2004; MCALLISTER and DYCK 2017). Furthermore, and despite the generally held idea that *w* mutants lack pigment because of defective transport of 3-hydroxy-kynurenine and 6-pyruvoyl tetrahydropterin into pigment granules (SULLIVAN *et al*. 1979; EVANS *et al*. 2008; GREEN *et al*. 2012; HERSH 2016; NAVROTSKAYA and OXENKRUG 2016), earlier studies had demonstrated physical absence of these organelles in the *w* mutants (NOLTE 1961; SHOUP 1966; NICKLA 1972). Could it be that all *Drosophila* mutants in the LRO-biogenesis pathway (LLOYD *et al*. 1998; KRAMER 2002; DELL'ANGELICA 2009; CHELI *et al*. 2010; HARRIS *et al*. 2011), including *w*, lacked zinc storage? To test this idea, total body zinc content was determined in the corresponding mutants.

### LRO-biogenesis mutants lack body zinc stores

Metal measurements by AAS and ICP-OES produced a good correlation between the two sets of data (Table 1). The analyzed genotypes readily separated into three groups according to their zinc content (Figure 2A). Null mutants in the *w* gene were all low in zinc; the reduction in metal content was specific for zinc and not seen for iron, copper and manganese measured in parallel (Table 1 and Figure 2B). In contrast, a *w*^+^ strain derived after 20 consecutive generations of single-pair backcrossing into *w* showed normal zinc accumulation.

**Figure 2.**
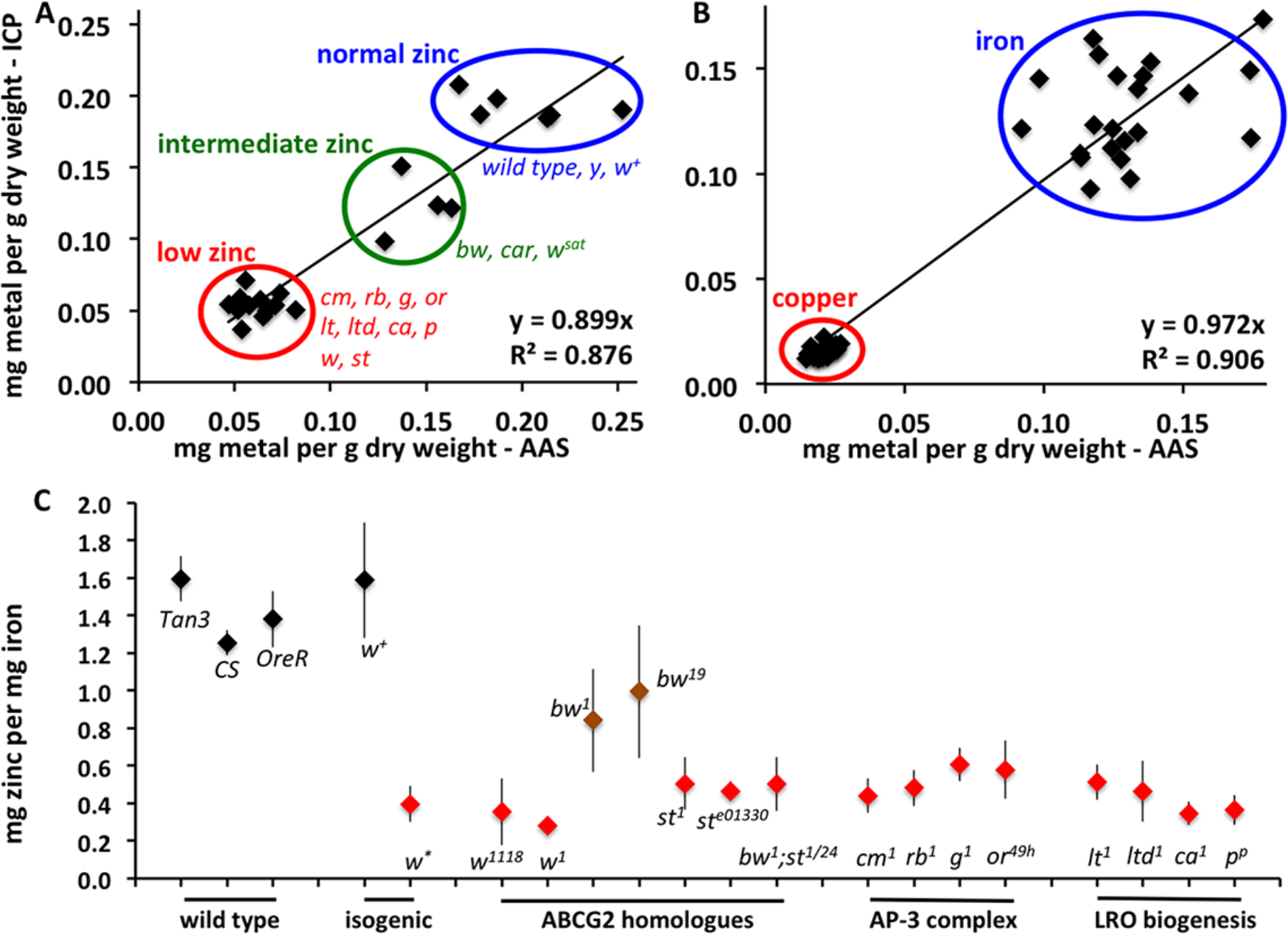
Mutants in the LRO biogenesis pathway have low body zinc content. (A) Linear regression between zinc determinations by AAS *versus* ICP-OES. The mean value determined for each genotype (Table 1), measured by both methods in biological replicates, is plotted. Stocks with normal, intermediate and low levels of zinc are readily identifiable. (B) Values for iron and copper are shown; here genotypes do not segregate. (C) Zinc-to-iron ratio was calculated for every independent measurement made (by either AAS or ICP-OES). Mean values and standard deviations are plotted.

We calculated the zinc-to-iron ratio from every measurement obtained per genotype irrespective of the technique used, and plotted the mean values and standard deviations from the mean (Figure 2C). Given the three-fold lower zinc-to-iron ratio in *w* mutants, we also tested *bw^1^* and *bw^19^*, which showed a minor reduction in zinc accumulation, whereas *st^1^* and *st^e01330^* were low in zinc, similar to *w* and to the double mutant *bw^1^*; *st^1/24^*. These results implicated the white-scarlet dimer (MACKENZIE *et al*. 2000) in *Drosophila* body zinc accumulation. Moreover, the AP-3 complex related mutants *cm^1^*, *g^1^*, *rb^1^* and *or^49h^* also had low body zinc as was true for the other LRO-biogenesis mutants *ltd^1^*, *ca^1^*, *p^p^* and *lt^1^*.

The response to dietary zinc chelation or supplementation was compared between *Tan3* wild type flies and *y^1^* mutants (used as an additional laboratory strain control) and *w*^*^ and *cm^1^* mutants (Figure 3). The control genotypes responded as expected to both treatments, reducing body zinc content when feeding on a diet supplemented with 200 μM N,N,N',N'-Tetrakis(2-pyridylmethyl)ethylenediamine (TPEN; a zinc-specific chelator) and increasing body zinc content on a diet supplemented with 1 mM zinc sulfate. In contrast, zinc chelation with TPEN had no effect on the body zinc content of *w*^*^ and *cm^1^* mutants, whereas zinc supplementation resulted in a small increase, barely reaching the body zinc content of *Tan3* wild type flies and *y^1^* mutants fed on 200 μM TPEN (Figure 3). These results suggest that *w* and *cm^1^* mutants are defective in zinc storage, lacking the part of wild type zinc that is chelatable with dietary TPEN.

**Figure 3.**
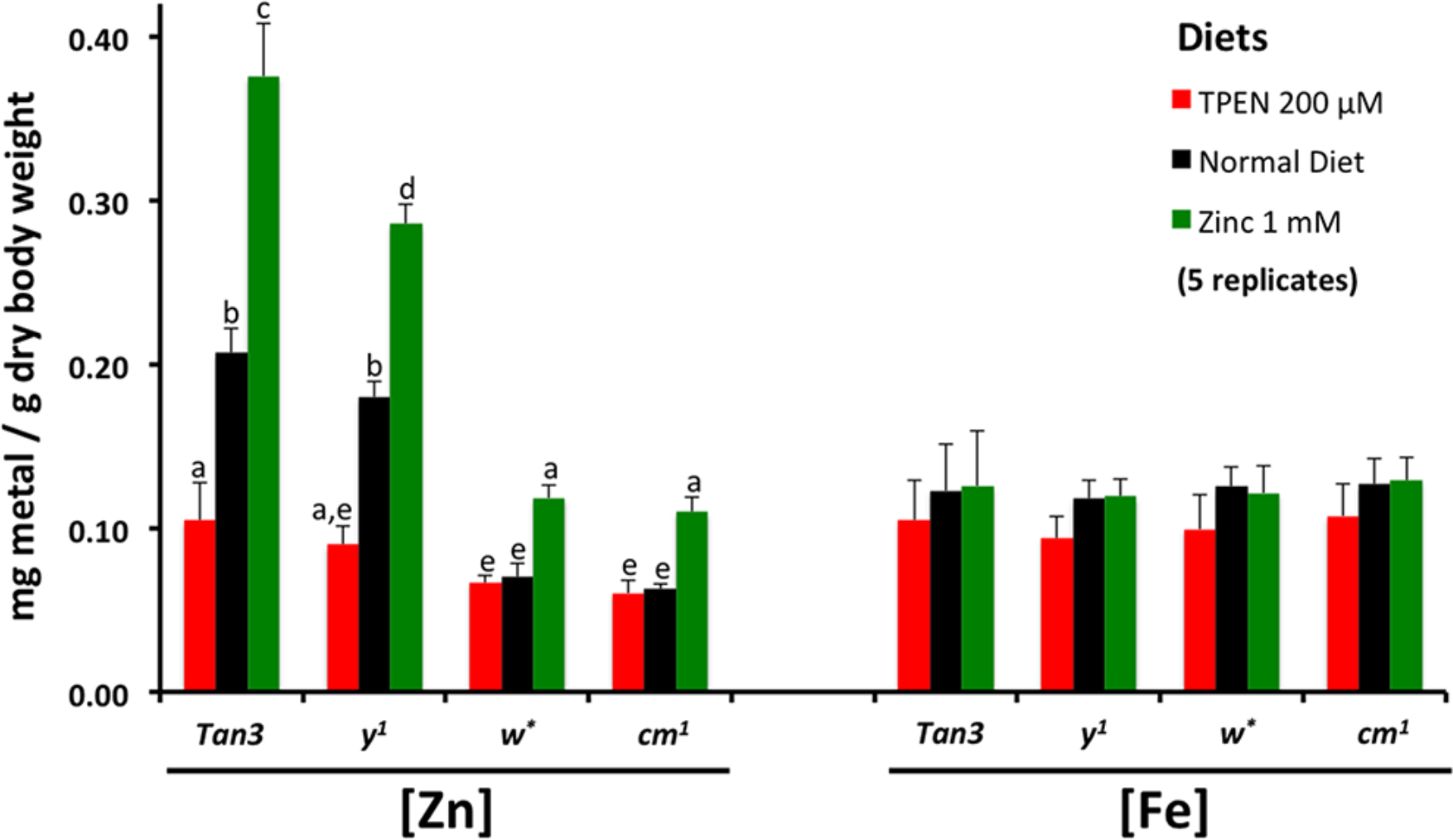
Zinc storage is affected in *w* and *cm* mutants. Two control strains (wild type *Tan3* and *y^1^*) and two low-zinc mutants (*w*^*^ and *cm^1^*) were grown on media with the zinc chelator TPEN (red bars) or supplemented with zinc sulfate (green bars) prior to measuring zinc and iron by AAS in whole flies from these populations. The control strains respond to the zinc treatments by changing body zinc stores, whereas this response is impaired in low-zinc mutants. Iron is unaffected. Two-way ANOVA showed differences by diet and by genotype; groups not different to each other from a *post-hoc* Tukey analysis are marked with the same lower-case letter.

### Malpighian tubule LROs are a major site for physiological zinc storage

A common feature of all LRO-biogenesis mutants is a reduction of pigment granules in their eyes. Null mutants in the *w* gene completely lack these organelles (NOLTE 1961; SHOUP 1966). Eye pigment granules are commonly rescued with the *mini-white* transgene (PIRROTTA *et al*. 1985), but the resulting stocks often remain low in body zinc (BETTEDI *et al*. 2011; GUTIERREZ *et al*. 2013). Thus, eye pigment cells are an unlikely location for the LROs mediating body zinc storage. Previous reports have documented zinc storage granules in the Malpighian tubules of *Drosophila hydei* (ZIEROLD and WESSING 1990) and of *Tumulitermes tumuli*, a termite species (STEWART *et al*. 2011). Moreover, another common feature of all LRO-biogenesis mutants is a reduction of riboflavin containing pigment granules in their Malpighian tubules (BREHME and DEMEREC 1942; NICKLA 1972). Generally, yellow Malpighian tubules correlated with normal zinc content, whereas loss of coloration correlated with low body zinc (Figure 4). Note, however, that *bw* and *st* mutants did not follow this rule, an exception to which we shall return later. For the remaining mutants presented in this study, our hypothesis was that disruption of LROs in the Malpighian tubules (as evidenced by loss of riboflavin granules) resulted in loss of zinc storage granules.

**Figure 4.**
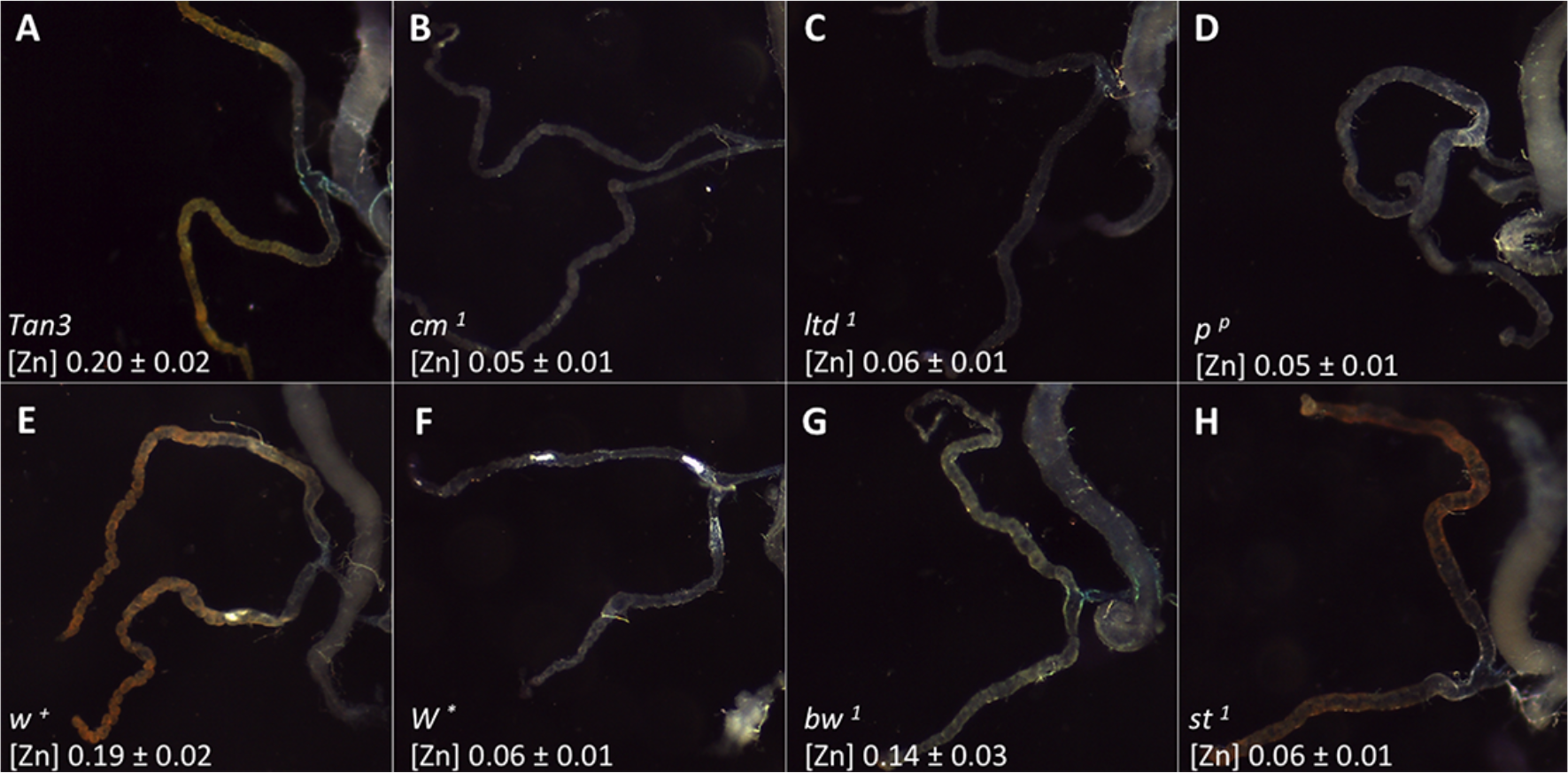
Mutants in the LRO biogenesis pathway fail to accumulate riboflavin in the Malpighian tubules. The posterior tubules of adult females are shown for the indicated genotypes along with the concentration of total body zinc in mg per g dry weight. (A) Riboflavin in wild type Malpighian tubules gives them their characteristic yellow-orange color (NICKLA 1972). (B-D) Three representative LRO biogenesis mutants are all colorless and low in zinc. A comparison between isogenic (E) *w*^+^ and (F) *w*^*^ suggests that the *w* gene is required for the accumulation of both riboflavin and zinc. (G) The *bw* mutants are severely reduced in their coloration and less so in their zinc content. (H) The *st* mutants show riboflavin coloration, but severely reduced zinc concentration.

To directly visualize zinc, synchrotron X-ray fluorescence imaging was performed (KORBAS *et al*. 2008; POPESCU *et al*. 2009; BOURASSA *et al*. 2014; JONES *et al*. 2015). Zinc was the major metal element detected in Malpighian tubules from *w*^+^ female flies (Figure 5A). In contrast, only trace amounts of zinc were detectable in Malpighian tubules from *w*^*^ flies (Figure 5B). Variation in the spectral emissions from other elements was minimal between the two samples, suggesting once again that the *w* gene affects specifically zinc accumulation in this tissue.

**Figure 5.**
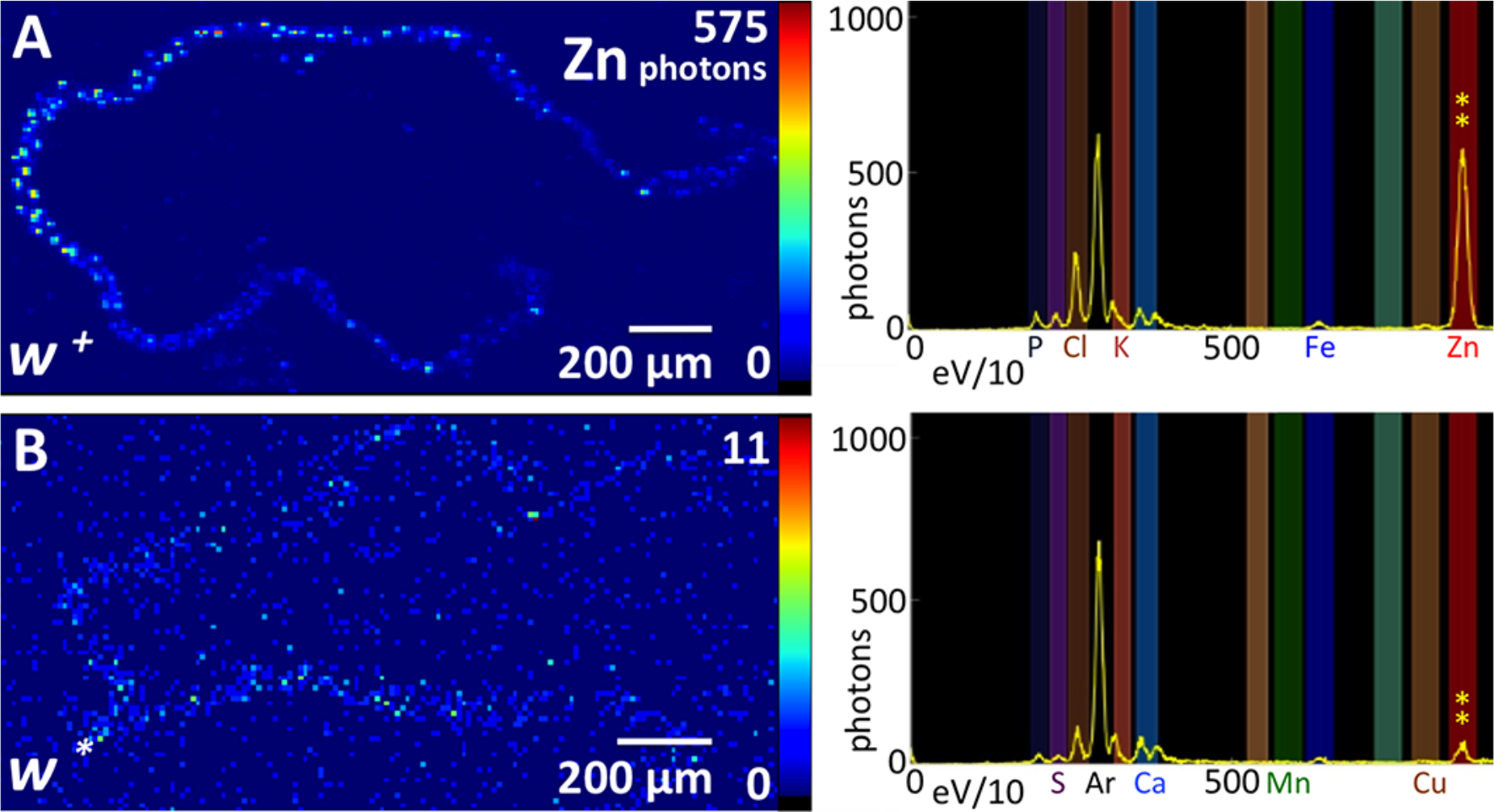
Synchrotron X-ray fluorescence microscopy demonstrates the presence of zinc in Malpighian tubules from adult female flies. Pixel resolution was 10 μm^2^. (A) Representative heat-map image for the zinc signal is shown for *w*^+^ Maplighian tubules. The spectral maximum for zinc emission is 575 photons per 100 ms (also indicated by the two yellow asterisks along with the full spectrum on the right panel). (B) Almost no zinc is detectable in Malpighian tubules from the *w* mutant.

To visualize zinc storage granules, Malpighian tubules were incubated with the zinc indicator Fluozin-3^AM^ and examined by confocal microscopy. Whereas the Malpighian tubules from adult female *w*^*^ flies showed only diffuse background signal (Figure 6A), in Malpighian tubules from adult female *w*^+^ flies multiple, distinct accumulations of fluorescence with a diameter of approximately 1 μm were observed (Figure 6B). These fluorescent structures, which we suggest are zinc storage granules, were present only in primary tubule cells and not in the supporting stellate cells (HALBERG *et al*. 2015).

**Figure 6.**
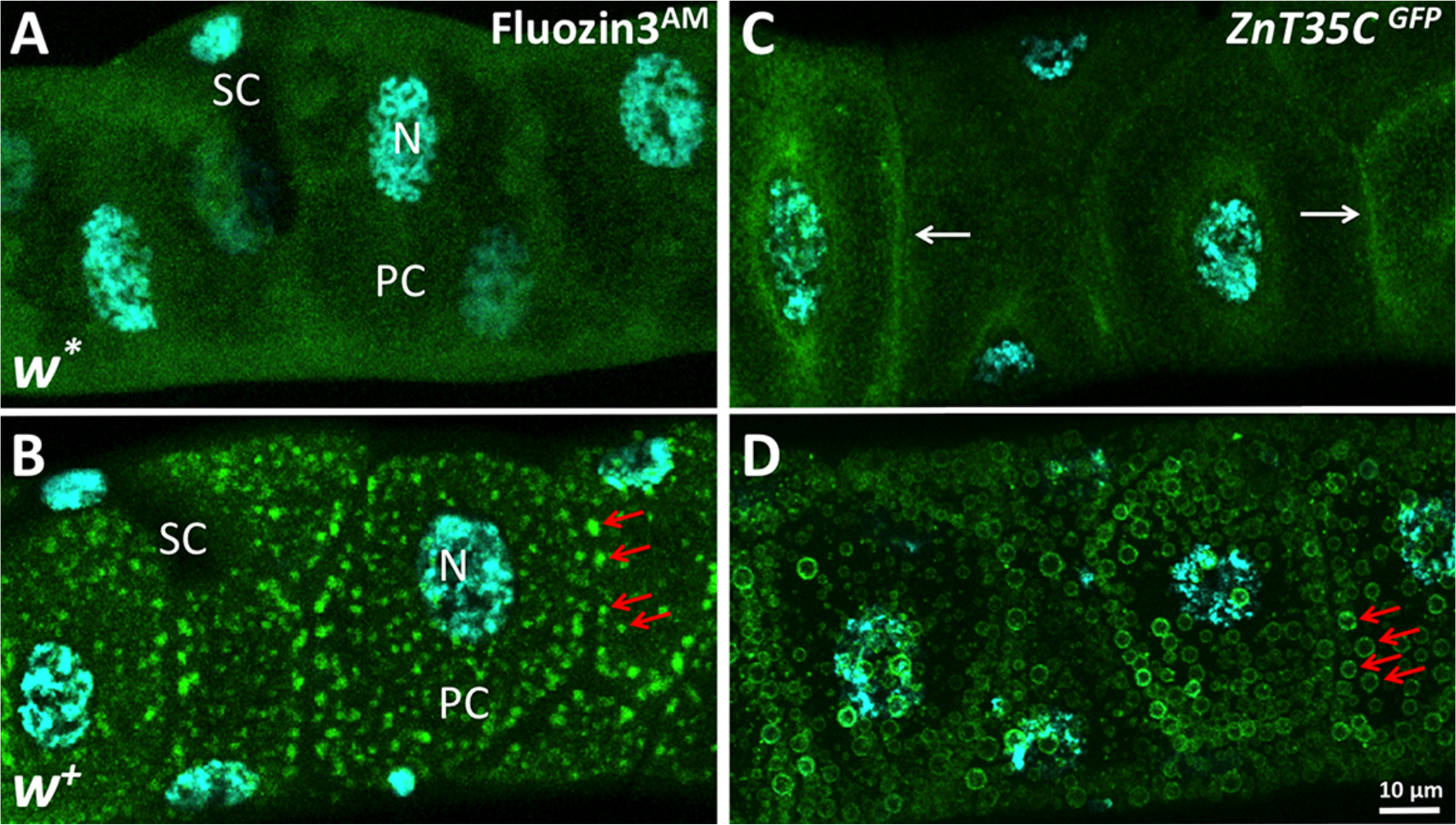
Zinc storage granules are located in the Malphighian tubules. Confocal images of Fluozin3^AM^ fluorescence, indicative of labile zinc in the Malpighian tubule of (A) a *w*^*^ female adult or (B) an isogenic *w*^+^ female adult. Red arrows point to a subset of zinc storage granules. (C) The protein trap line *y,w; Znt35C^GFP^* was used to monitor the subcellular localization of ZnT35C, confirming previous observations that it associates with the plasma membrane (white arrows). (D) The same reporter in the *w*^+^ background clearly marks the zinc storage granule membrane (red arrows). All flies were grown on a diet supplemented with 5 mM zinc sulfate. SC – stellate cell; PC – primary cell; N – nucleus.

Zinc accumulation in vesicles normally depends on specialized transporters. ZnT8 for example, is responsible for zinc entry into pancreatic insulin-granules (POUND *et al*. 2009), while ZnT3 takes over this function in glutamatergic vesicles of the mossy fibers (COLE *et al*. 1999; MCALLISTER and DYCK 2017). We hypothesized that the best candidate *Drosophila* zinc transporter to mediate this function was ZnT35C, which is phylogenetically related to human ZnT3 and ZnT8 (LYE *et al*. 2012) and exclusively expressed in the Malpighian tubules (YEPISKOPOSYAN *et al*. 2006; CHI *et al*. 2015; YIN *et al*. 2017). We used a strain that inserts GFP into the endogenous ZnT35C open reading frame (NAGARKAR-JAISWAL *et al*.2015) and observed the subcellular localization of the tagged transporter in Malpighian tubules of adult female flies grown on a zinc-supplemented diet. In the *w* mutant that lacks LROs, ZnT35C was localized in the proximity of the plasma membrane (Figure 6C), consistent with previous observations in Malpighian tubules from the larvae (YEPISKOPOSYAN *et al*. 2006; YIN *et al*. 2017). In contrast, ZnT35C^GFP^ clearly marked a subset of LROs in *w*^+^ flies, which correspond to zinc storage granules (Figure 6D).

## DISCUSSION

### Zinc storage in *Drosophila melanogaster* and other animals

We propose that the zinc storage granules in primary cells of the *Drosophila* Malpighian tubules (Figure 6) have a similar function to ferritin-containing Golgi-related vesicles in iron cells of the middle midgut (LOCKE and LEUNG 1984; MISSIRLIS *et al*. 2007). The latter serve for iron storage as a physiological parallel of liver ferritin (MEHTA *et al*. 2009), and the former do the same for zinc storage. Zinc storage granules, first described in *Drosophila hydei* (ZIEROLD and WESSING 1990), have also been observed in termites (STEWART *et al*. 2011). Twenty-four species of flies (from the Drosophilidae and the Tephritidae families) have similar zinc content (SADRAIE and MISSIRLIS 2011; REMPOULAKIS *et al*. 2014), suggesting that the function of Malpighian tubules in zinc storage and excretion (CHI *et al*. 2015; YIN *et al*. 2017), is evolutionarily conserved (HALBERG *et al*. 2015).

As there is no generally accepted site for body zinc storage in humans or other mammals, it is worth investigating whether zinc-containing granules such as those present in the pancreas (SCOTT and FISHER 1938; TIMM and NETH 1958; KAWANISHI 1966; RUTTER *et al*. 2016; MARET 2017) or in intestinal paneth cells (OKAMOTO 1942; GIBLIN *et al*. 2006) serve as a reservoir for this metal, as may be the case, alternatively or additionally, for bone depositions of zinc (BERG and KOLLMER 1988; HUANG *et al*. 2007). The only animal where a zinc storage site has been proposed is the nematode *Ceanorhabditis elegans* (ROH *et al*. 2012; WARNHOFF *et al*. 2017). Upon feeding on a diet with high content of zinc, the *C. elegans* generates new vesicles within intestinal cells to store the excess metal. Thus, zinc storage granules, or zincosomes as they have been also called (BEYERSMANN and HAASE 2001; COLVIN *et al*. 2016), appear to be a conserved LRO in animal biology.

### Different types of LROs defined by *bw* and *st*

The Malpighian tubules are also known to store riboflavin into LROs (NICKLA 1972). Our results suggest that the brown-white dimer is primarily required for the formation of the riboflavin LRO (Figure 4G), whereas the scarlet-white dimer is required for the zinc storage granule (Figure 4H). All other genes tested and known to be involved in the biogenesis of LROs affected both riboflavin accumulation and zinc storage (Figure 4). This poses an interesting cell biology question, as the function of the Rab32, AP3, HOPS and BLOC complexes is understood as enabling the trafficking (segregation) of transporters such as ZnT35C to the LRO, defining in this way the organelle’s identity (LLOYD *et al*. 1998; DELL'ANGELICA *et al*. 2000; BULTEMA *et al*. 2012; GERONDOPOULOS *et al*. 2012; BONIFACINO and NEEFJES 2017). Our description of two types of LROs in the Malpighian tubule primary cells (Figure 7) requires an explanation of how the brown-white dimer is segregated away from the scarlet-white dimer to give rise to different types of LROs.

**Figure 7.**
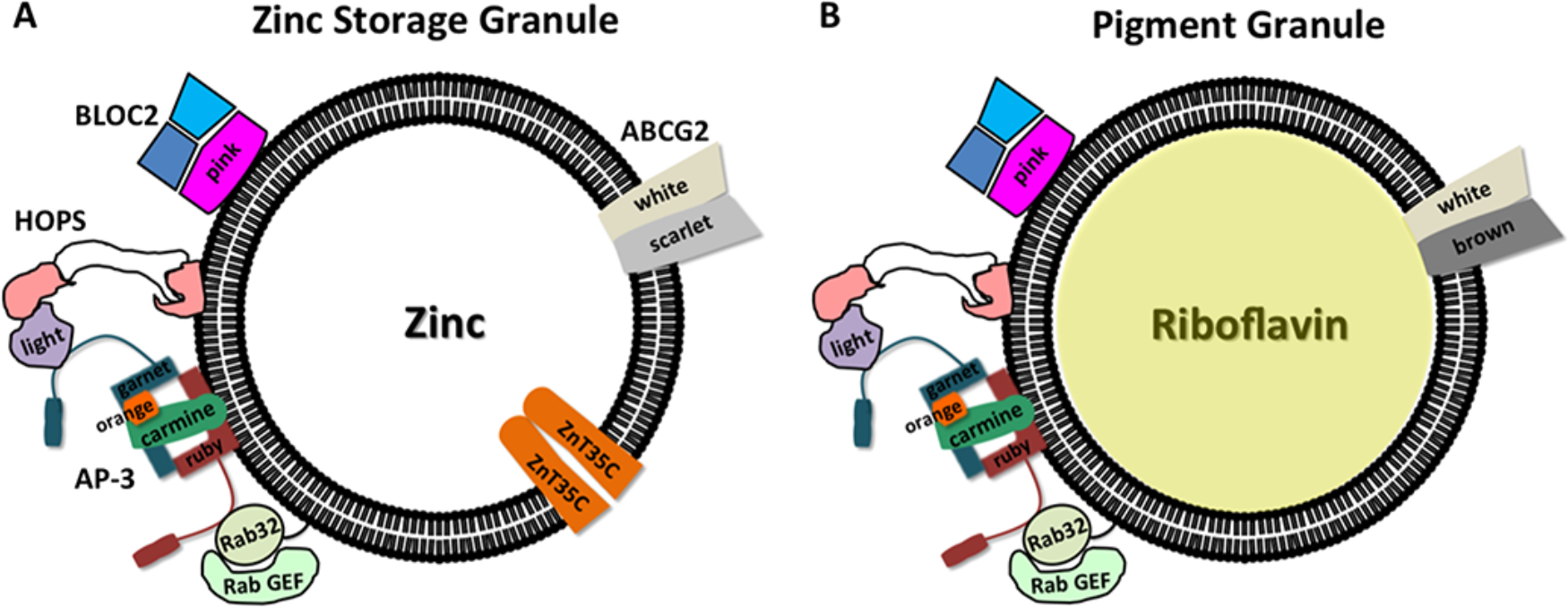
Schematic representation of the (A) zinc storage granule and (B) riboflavin pigment granule, in primary cells of the Malpighian tubules. On the left of each vesicle known players in protein trafficking are shown, common for the biogenesis of both types of LROs. Transporters are on the right side. It is unclear how cells differentially give rise to the two LROs.

### On the function of the ABCG2 transporters

The human ABCG2 has been studied extensively, because of early reports associating ABCG2 overexpression with the development of cancer resistance to drugs inhibiting the cell cycle by intercalating into DNA (CHEN *et al*. 1990). The consensus for the mechanism of action of ABCG2 is its direct activity in exporting drugs from cells, however the apparent lack of specificity in the transported molecules is puzzling: more than 200 substrates have been described for ABCG2 (HORSEY *et al*. 2016). Amongst these substrates, one is riboflavin (VAN HERWAARDEN *et al*. 2007), another is cGMP (DE WOLF *et al*. 2007; EVANS *et al*. 2008). Does the function of ABCG2 relate more to the failure of formation of a LRO (LLOYD *et al*. 2002) in the cells or animal models where it has been silenced, and less to the specific transport of the various substrates it has been claimed to move across membranes? At least in *Drosophila*, the *w* gene is required for the process of pigment granule formation *per se*.

### On the use of the *w* mutant as a control in *Drosophila* experiments

Many authors warn against the possible alterations of normal cell physiology in the *w* mutant and consider implications of using it as the major control strain in experiments (CAMPBELL and NASH 2001; BORYCZ *et al*. 2008; CHETVERINA *et al*. 2008; KRSTIC *et al*. 2013; CHAN *et al*. 2014; XIAO and ROBERTSON 2016). Direct implications for the field of *Drosophila* zinc biology, almost entirely based on transgenes carrying the *mini-white* marker, have been raised before (AFSHAR *et al*. 2013; RICHARDS and BURKE 2016).

### A potential role for zinc in human Hermansky-Pudlak syndrome

Mutations in 9 different genes can cause Hermansky-Pudlak syndrome in humans Hermansky-Pudlak syndrome is characterized by oculocutaneous albinism, a platelet storage pool deficiency and lysosomal accumulation of ceroid lipofuscin (SEWARD and GAHL 2013). Patients with the genotypes HPS-1, HPS-2, or HPS-4, are predisposed to interstitial lung disease and may develop granulomatous colitis. Hypopigmentation is the prominent feature of HPS, attributable to the disrupted biogenesis of LROs (WEI *et al*. 2013). Is zinc homeostasis altered in human patients diagnosed with Hermansky-Pudlak syndrome? To our knowledge this question has not been addressed. We conclude by pointing out that fly mutants lacking zinc storage granules correspond to known genetic alterations in Hermansky-Pudlak patients. Besides the identification of *p* as the homologue of the Hermasky-Pudlak syndrome type 5 gene (FALCON-PEREZ *et al*. 2007; SYRZYCKA *et al*. 2007), *rb* encodes for the type 2 syndrome protein (GOCHUICO *et al*. 2012) (a second neuronal-specific human homologue of the β subunit of AP-3 was recently related to an early-onset epileptic encephalopathy with optic atrophy (ASSOUM *et al*. 2016)), and *g* encodes for the δ subunit of the AP-3 complex, recently shown to define a new type of the Hermansky-Pudlak syndrome (AMMANN *et al*. 2016). Likewise, the corresponding mouse mutants are known to lack zinc granules (KANTHETI *et al*. 1998). Thus, the regulation of zinc homeostasis and trafficking in Hermansky-Pudlak patients deserves further investigation.

## AUTHOR CONTRIBUTIONS SECTION

FM mapped the *poco-zinc* mutant. CTG was directly involved in all other experiments. ARA collaborated in tissue chemistry/confocal microscopy and, together with TK and SW, performed the X-ray fluorescence imaging. MB, BO and FM helped with metal determinations. FM directed the work and wrote the paper, which was reviewed, corrected and endorsed by all authors.

## ACKNOWLEDGMENTS

The authors thank the Bloomington Drosophila Stock Center for the flies used in this study. We acknowledge Refugio Rodríguez Vázquez for providing access to the atomic absorption spectrometer, Alma Isabel Santos Díaz for participating in experiments as part of her social service training in the laboratory, Marcos Nahmad, Irene Miguel-Aliaga, and John F. Allen for critical comments on the manuscript. Use of the Stanford Synchrotron Radiation Lightsource, SLAC National Accelerator Laboratory, is supported by the U.S. Department of Energy, Office of Science, Office of Basic Energy Sciences under Contract No. DE-AC02-76SF00515. The SSRL Structural Molecular Biology Program is supported by the DOE Office of Biological and Environmental Research, and by the National Institutes of Health, National Institute of General Medical Sciences (including P41GM103393). The authors thank Nicholas P. Edwards and Courtney M. Krest for excellent beam line support and José Mustre de León for approval of travel expenses. The MARS6 microwave digestion system and the PerkinElmer Optima™ 8300 ICP-OES instrument were acquired thanks to the CONACYT infrastructure grant #268296. CONACYT also supported Carlos Tejeda-Guzmán & Abraham Rosas-Arellano with Ph.D. (#299627) and postdoctoral (#189290) fellowships, respectively.

## Footnotes

1 These authors contributed equally to this work.

